# Primary Cilium Disassembly in Mammalian Cells Occurs Predominantly by Whole-Cilium Shedding

**DOI:** 10.1101/433144

**Authors:** Mary Mirvis, Kathleen Siemers, W. James Nelson, Tim Stearns

**Affiliations:** Departments of Molecular and Cellular Physiology, Stanford University, Stanford, CA, 94305; Departments of Molecular and Cellular Physiology, Biology Stanford University, Stanford, CA, 94305; Departments of Molecular and Cellular Physiology, and Biology, and Genetics, Stanford University, Stanford, CA, 94305

## Abstract

The primary cilium is a central signaling hub in cell proliferation and differentiation, and is built and disassembled every cell cycle in most animal cells. Disassembly is critically important: misregulation or delay of disassembly leads to cell cycle defects. The physical means by which cilia are disassembled are poorly understood, and thought to involve resorption of disassembled components into the cell body. To investigate cilium disassembly in mammalian cells, we used rapid live-cell imaging to comprehensively characterize individual disassembly events. The predominant mode of disassembly was rapid cilium loss via deciliation, in which the membrane and axoneme of the cilium was shed from the cell. Gradual resorption was also observed, as well as events in which a period of gradual resorption ended with rapid deciliation. Deciliation resulted in intact shed cilia that could be recovered from culture medium and contained both membrane and axoneme proteins. We modulated levels of katanin and intracellular calcium, two putative regulators of deciliation, and found that excess katanin promotes disassembly by deciliation, independently of calcium. Together, these results demonstrate that mammalian ciliary disassembly involves a tunable decision between deciliation and resorption.

## Introduction

Cilia are present throughout all branches of the eukarya, and are adapted to facilitate interactions between cells and their surrounding environment through the transduction of molecular and mechanical signals, and the ability to generate motion (1–3). All cilia share a core architecture consisting of a basal body (centriole), a core of stable microtubule doublets (axoneme), and a distinct ciliary membrane (4–6). In many cases, including vertebrate primary cilia and *Chlamydomonas* motile cilia, the presence of a cilium is closely linked with the cell cycle, generally forming in interphase (G0) and disassembling prior to mitosis (6–12). Misregulation or delay in primary cilia disassembly leads to defects in cell cycle progression, which underlie aberrant developmental and homeostatic phenotypes in ciliopathies and many cancers (7,13–26).

Despite recent progress in the identification of molecular players and pathways regulating primary cilia disassembly (12,22,24,27–36), the physical mechanisms by which mammalian cilia are ultimately lost have not been identified. In *Chlamydomonas*, cilia have been reported to be resorbed into the cell body preceding cell division with rates of approximately 0.3-0.7 μm/min. (26,27,32,35–45). Alternatively, under conditions of stress or pharmacological induction, comparatively rapid removal of cilia, defined as the concurrent release of membrane and axoneme from the cell body, has been described in many species as deciliation, ciliary excision, shedding, deflagellation, or flagellar autotomy (46–53). Whether deciliation occurs as part of normal primary cilium behavior, and contributes to cell cycle-linked disassembly, is unknown. In this work, we use rapid long-term live cell imaging to determine whether either of these physical mechanisms (resorption or deciliation) contributes significantly to mammalian cell cycle-associated cilia loss. Ours is the first comprehensive study of mammalian primary cilium disassembly events to determine the dynamics and underlying physical means of ciliary loss.

## Results

### Ciliary structures in cells undergoing serum-induced cilium disassembly

To assess the physical processes underlying ciliary disassembly, we observed ciliary morphology in a population of mammalian cells entering the cell cycle and undergoing ciliary disassembly. We manipulated serum level in the culture medium to synchronize the disassembly of primary cilia (24,54) in IMCD3 cells expressing SSTR3::GFP (55), a fluorescent plasma membrane marker that is concentrated in the ciliary membrane (Fig. 1A, S1A). The majority of serum-starved cells (60 +/-9.09%) were ciliated. Serum stimulation for 6 h resulted in a decrease in the fraction of cells with a cilium (30 +/-0.4%) to levels comparable to asynchronously cycling cells (Fig. 1B). Serum-induced ciliary disassembly has been shown to require the function of HDAC6, a deacetylase of microtubules and cortactin [refs]. Cells treated with 2 μM tubacin, an HDAC6 inhibitor, failed to undergo serum-induced ciliary loss (Fig. 1B) (24,56,57). Mitotic cells accumulated following serum stimulation (Fig. SlB), but were inhibited or delayed in tubacin-treated cells (Fig. S1B), consistent with a requirement for ciliary disassembly prior to mitotic entry (10,11,58–60). Thus, serum stimulation induces synchronous, cell cycle-linked ciliary disassembly in these cells.

**Fig 1.**
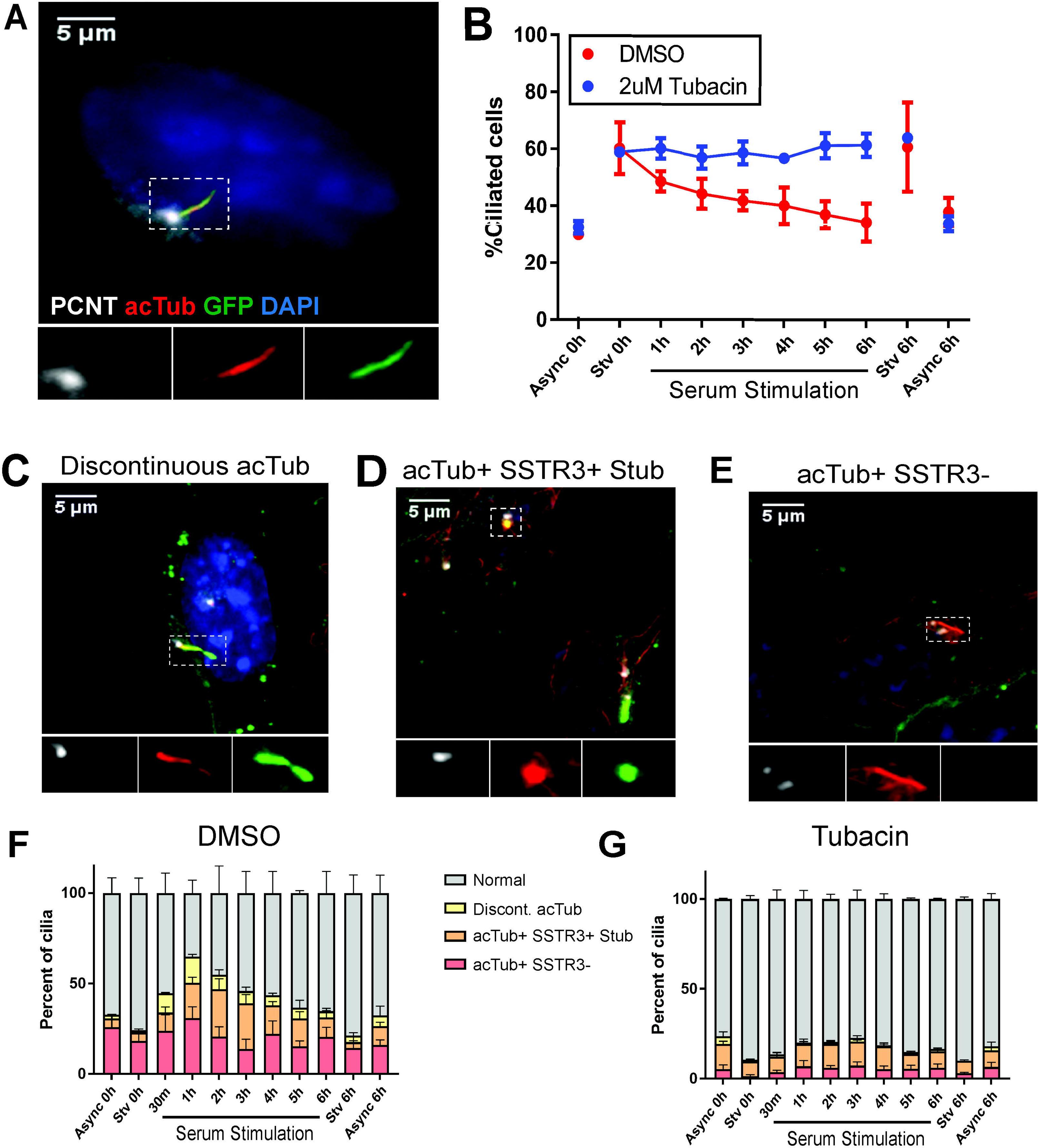
Serum stimulation of IMCD3 cells reveals non-canonical ciliary structures. Serum starved IMCD3-SSTR3::GFP cells were stimulated with 10% serum in order to synchronize ciliary disassembly. A, C-E) Cells were fixed at indicated time points after serum addition and immunostained for pericentrin to mark the basal body (PCNT, white) and acetylated tubulin to mark the axoneme (acTub, red). A) Morphology of a normal, intact cilium in a starved cell. B) The population of ciliated cells quantified over a serum stimulation time course. Asynchronous (Async) and serum starved (Stv) controls were included at 0 hr and 6 hr. Cells treated in parallel with 2 μM tubacin. C-E) Non-canonical ciliary structures identified in serum stimulated cell populations. C) Discontinuous acTub staining, in this case accompanied by narrowing of the membrane. D) A ciliary stub, marked by punctate acTub and SSTR3 fluorescence. E) Full-length axoneme marked by acTub, lacking corresponding SSTR3::GFP signal. F-G) Stacked plots of cilia morphologies observed during serum stimulation in (F) DMSO and (G) tubacin-treated cells. Quantifications are based on means of 3 independent experiments with 150-200 cells analyzed per condition per replicate. Error bars – S.E.M. Statistical significance was assessed by unpaired t-test, *p<0.05, **p<0.01, ***p<0.001.

Seeking a more systematic understanding of disassembly behaviors, we next characterized cilia morphology in cells undergoing ciliary disassembly during the 6-hr serum stimulation time course. We identified three types of cilium structures as likely disassembly intermediates in fixed cells expressing SSTR3::GFP (Fig. 1C-G): 1) a discontinuous axoneme, marked by a gap in acTub staining in all z-planes, (“Discontinuous acTub”, Fig. 1C); 2) a ciliary stub, characterized by a short (<1 μm) cilium positive for both acTub and SSTR3 membrane fluorescence (“acTub+ SSTR3+ Stub”, Fig. 1D); and 3) an axoneme marked by linear acTub fluorescence (>1 μm), but lacking corresponding membrane SSTR3 fluorescence (“acTub+ SSTR3-”, Fig. 1E). All intermediate structures were rare in serum-starved cells, but at 2-3 h after serum-stimulation they comprised the majority of detectable cilia (Fig. 1F). Tubacin treatment prevented the enrichment of all three structures in serum-stimulated cells (Fig. 1G), indicating that these structures are most likely representative of disassembling cilia. These results suggest that the mechanism of ciliary disassembly might be variable even within a single cell population, and we next sought to use live cell imaging to observe the dynamics of cilia disassembly.

### Rapid deciliation is the predominant mode of ciliary disassembly

To observe the spatio-temporal features of cilium disassembly directly, we generated IMCD3-SSTR3::GFP cells which stably co-expressed a basal body marker (mCherry-PACT) (see Materials & Methods). Live cells were imaged immediately following serum stimulation with full confocal stacks acquired continuously for 6-12 h at 90 sec intervals. Apically-facing cilia were selected for analysis, to avoid artifacts caused by deformations of basally-facing cilia by the glass imaging dish. Observations of serum-starved cells were used as controls for ciliary behaviors not associated with disassembly (Fig. 2A.1, Fig. S2A).

**Fig 2.**
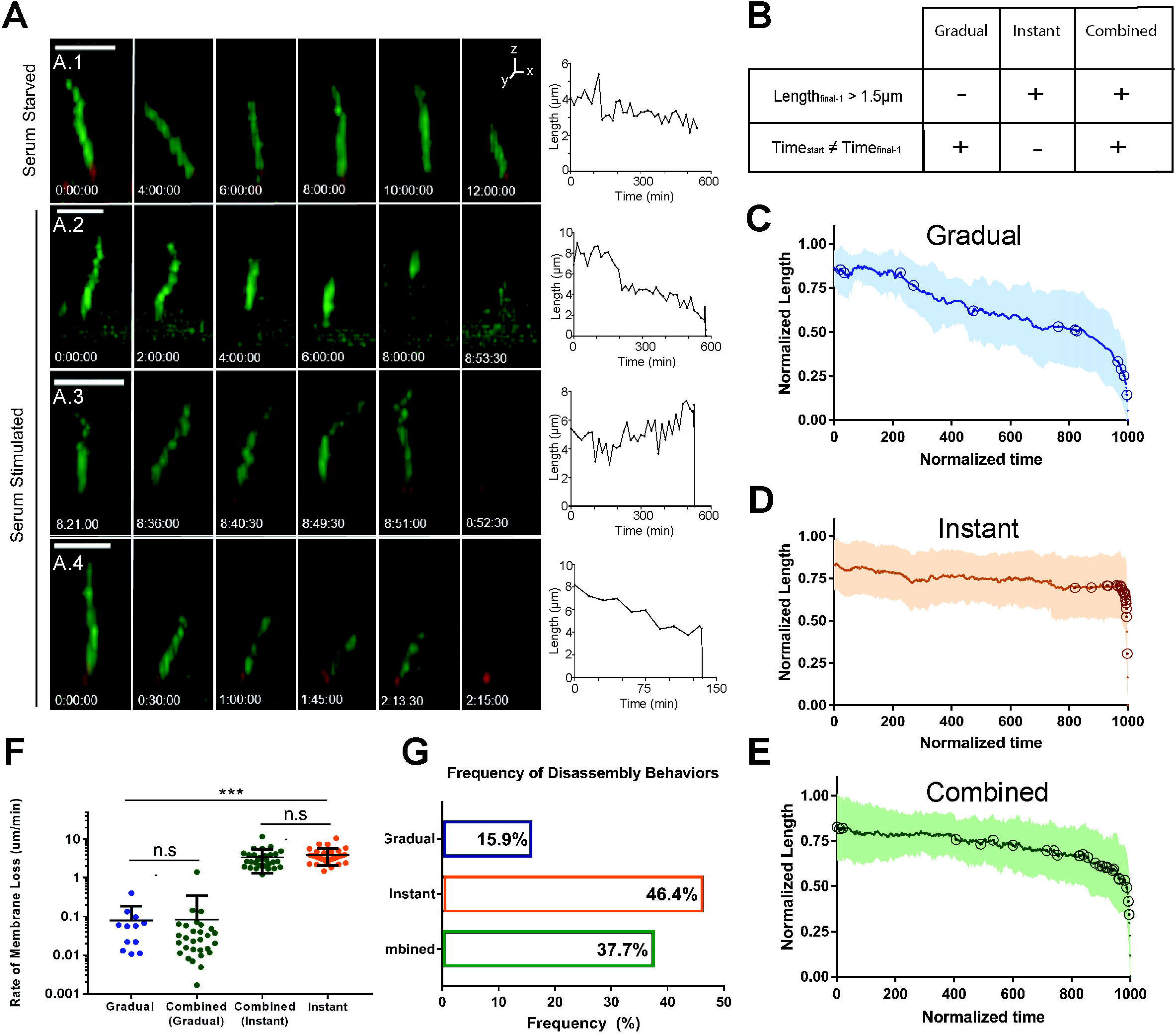
Live-cell analysis of cilia disassembly reveal highly heterogeneous dynamics. A) Still images and raw length curves representing dynamics of individual cilia disassembly events. Length measurements based on SSTR3::GFP fluorescence. Scale bar 5 μm. A.1) Control serum starved cilium measured over 12 hr period undergoes slight length change of 1.66 μm at a rate of 0.003 μm/min. A.2) Gradual disassembly with a rate of 0.016 μm/min. A.3) Instant disassembly, with an approximate minimum rate of 4.72 μm/min. A.4) Combined disassembly, consisting of an initial stage of gradual shortening at a rate of 0.029 μm/min, followed by instant loss at a minimum rate of 5.69 μm/min. B-E) Semi-automated analysis of disassembly events. B) Criteria for defining gradual, instant, and combined behaviors. See text and Fig. S2 for more details. C-E) Cumulative normalized length vs. time curves for all (C) *gradual*, (D) *instant*, and (E) *combined* disassembly events. Open circles indicate individual event start points. F) Disassembly rates within dynamic groups. Significance was determined by unpaired t-test, *p<0.05, **p<0.01, ***p<0.001. Error bars – S.D. H) Relative frequency of disassembly behaviors (n=70).

Video sequences of disassembling cilia revealed a striking range of disassembly behaviors (Fig. 2A and Movies S1-4). We grouped these behaviors into three categories: *gradual* -cilium length reduction over at least two consecutive time points resulting in terminal cilium loss (e.g., Fig. 2A.2, 0.02 μm/min; T_start_ ≠ T_final-1_, see Fig. S2); *instant* -a single discrete cilium loss event within a single imaging frame, i.e. 30-90 sec (e.g., Fig. 2A.3, ≥4.72 μm/min L_final-1_ > 1.5 μm); and *combined* – a period of *gradual* disassembly directly followed by *instant* loss (e.g., Fig. 2A.4, gradual phase (0.03 μm/min;), preceding rapid loss within 46 seconds, ≥5.69 μm/min).. To reduce bias in our cumulative analysis, we developed an algorithm to normalize ciliary length fluctuations to controls, identify a disassembly start point, and assign each event to one of the three categories described above (Fig. 2B & S2, Materials & Methods). Disassembly curves were normalized by time (to 1000 arbitrary units) and ciliary length (to the maximum length of each cilium) (Fig. 2C-E). The *gradual* averaged curve shows early ciliary shortening with event start points distributed along the curve, followed by a period of consistent shortening in the last ~150 normalized time units as the slope of the curve increases (Fig. 2C). The *instant* averaged curve appears nearly horizontal until the last point; start points were nearly all clustered in the last ~10 time units (Fig. 2D). The *combined* averaged curve features a period of slight slope characteristic of *gradual* dynamics followed by an *instant* cilia loss event, with start points distributed along the curve (Fig.2E).

Ciliary disassembly rates spanned several orders of magnitude (10^−3^ – 10^1^ μm/min (Fig.2F)), demonstrating the dynamic heterogeneity of the process. The slowest *gradual* disassembly events were on the order of several hours, while the fastest *instant* events (10^1^ μm/min) represented behaviors in which an entire cilium was lost in less than 90 seconds. Within the *combined* category, the rate of the first *gradual* step (0.08 μm/min ± 0.26) and second *instant* step (3.41 μm/min ± 2.10) were not significantly different from those of the *gradual*-only (0.08 μm/min ± 0.11) and *instant*-only (3.88 μm/min ± 1.80) rates, respectively (Fig. 2F). Further, the majority of ciliary length (72.6 ± 4.5%) was lost during the *instant* stage. Thus, *combined* disassembly likely represents both *gradual* and *instant* mechanisms occurring within one cilium disassembly event, rather than an independent third behavior with biphasic dynamics.

Strikingly, we found that the *instant* and *combined* groups together comprised 83.1% (n=70) of the observed disassembly events (Fig. 2G). Thus, events with *instant* dynamics, which occur within seconds, are the predominant behavior for terminal ciliary loss. Rates of *gradual* disassembly were heterogeneous, but roughly consistent with previously reported rates of resorption (36,43,44), while *instant* dynamics were significantly more rapid than would be expected for resorption. To test whether *instant* disassembly dynamics might be consistent with deciliation, we examined the dynamics of deciliation induced by dibucaine, which likely raises intracellular calcium (47,49,50,61). Dibucaine-induced deciliation had qualitative features and a dynamic profile similar to serum-induced *instant* disassembly (Fig. S3 & 2A.2). Therefore, we hypothesized that deciliation is the physical mechanism underlying *instant* cilia loss.

In a rare instance, we observed direct shedding of the entire visible cilium from the surface of a serum-stimulated cell (Fig. 3A, Movie S5). The shed cilium in the z-stack shown in Fig. 3A has a fragmented appearance. Due to the rapid time scale of the shedding event, the shed cilium can travel a considerable distance (>1μm) in the time between individual z-slices (roughly 2.6 sec) within the stack. As a result, the shift in apparent location of the same object in subsequent slices causes the final imaged object to appear distorted – artificially elongated or with the appearance of separated fragments. We note that the segments in such sequences were visualized as a co-migrating cluster, rather than dispersing independently, consistent with the interpretation that they are parts of one structure imaged at slightly different times. Therefore, we interpret the fragmented appearance of the cilium in Fig. 3A as the result of motion of the imaged object during acquisition, rather than a biological change in morphology, and hypothesize that these images represent a deciliation event.

**Fig 3.**
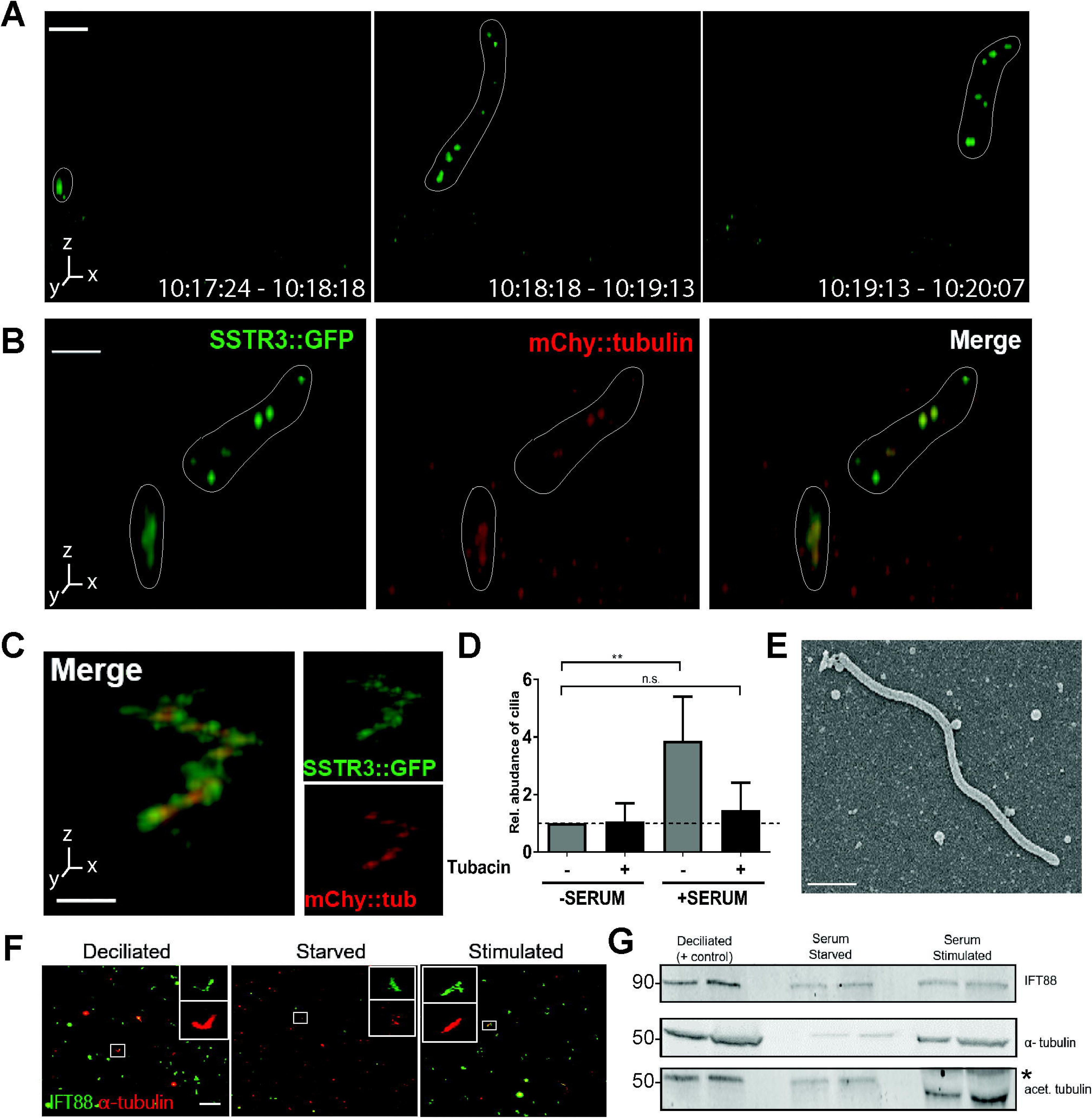
Observation and validation of whole-cilium shedding. A) Serum stimulated cell, imaged at 54-second intervals. A short ciliary membrane (in white outline) is visibly shed from the cell surface. Time labels are interval of stack-acquisition. bar, 5 μm. B) Partial ciliary shedding in IMCD3-SSTR3::GFP cells transiently expressing mCherry-α-tubulin, imaged at 30 s-intervals. Intact and shed portions of the cilium are demarcated with a white outline. Tubulin is visible in shed fragments. Scale bar, 3 μm. C-E) Ciliary immune-capture. Serum stimulation media were incubated on immobilized anti-SSTR3 antibody. Isolated cilia were observed by C-D) confocal fluorescent microscopy (scale bar, 2 μm) and E) SEM. Scale bar, 500nm. C-D) Immune-captured cilium from IMCD3-SSTR3::GFP cells transfected with mCherry-α-tubulin. D) Normalized quantification of (C), defined as elongated SSTR3+ objects at least 1.5 μm in length. Tubacin treatment reduces the prevalence of captured cilia. F-G) Filter-spin concentration of shed cilia. Serum stimulation media from IMCD3 cells with no endogenous fluorescence was concentrated either by (F) vacuum & centrifugation filtration for immunofluorescence or (G) centrifugation pelleting for subsequent Western blot. Media from artificially deciliated cells were included as a positive control. F) Filtration-concentrated samples were processed for immunofluorescence against IFT88 and α-tubulin. Scale bar, 20 μm. Representative insets show full-length cilia: 6.82μm (deciliated), 5.94μm (starved), and 6.10μm (stimulated). G) Western blot of concentrated material shows presence of ciliary markers at expected molecular weights: α-tubulin & acetylated tubulin. Molecular weights are indicated (kDa). Asterisk (*) denotes position of a large BSA band present in samples derived from serum-containing medium. Two replicates per condition are shown as side-by-side bands.

The rapid nature of the deciliation event shown in Fig. 3A (<54 secs), and the diffusion of the shed ciliary remnant(s) away from the site of origin is consistent with the *instant* loss dynamics found in 83.1% of serum-induced disassembly events (Fig. 2G) and in dibucaine-treated cells (Fig. S3). In addition, cilia shed under our culture conditions from cells expressing mCherry-α-tubulin contained tubulin in shed ciliary fragments (Fig. 3B) suggesting that the axoneme is shed together with the ciliary membrane. These results further support the hypothesis that *instant* cilium loss represents ciliary disassembly via deciliation.

### Recovery of whole cilia from culture media demonstrates that deciliation occurs during ciliary disassembly

If deciliation is a major mode of cilium disassembly, we expect to be able to recover whole cilia from culture medium. These cilia should be of plausible length for whole cilia (at least 1.5μm, see Fig. S2), and contain membrane- and axoneme-specific components. We developed two methods to enrich for ciliary fragments spontaneously released at low concentration from serum-stimulated cells (Fig. 3C-G, S4). First, we used immuno-capture of fluorescently-labeled cilia to directly visualize unperturbed ciliary fragments. Culture medium from serum-stimulated IMCD3::SSTR3-GFP cells transiently expressing mCherry-α-tubulin was incubated on imaging dishes coated with an antibody against the extracellular domain of SSTR3 (Fig. S4A). Samples were imaged, without fixation, by fluorescence; we were indeed able to reproducibly identify cilia marked by both SSTR3-GFP and mCherry-α-tubulin captured in this manner (Fig. 3C). We interpret that the jagged appearance of the cilia is due to thermal motion of the sample during stack acquisition of material that is not firmly fixed to the substrate. Compared to control serum-starved medium, serum-stimulated medium yielded a 4-fold increase in captured cilia. In addition, pre-treatment with tubacin decreased the number of captured cilia to control levels (Fig. 3D). Finally, we found objects with dimensions consistent with intact shed cilia after fixation and imaging by scanning electron microscopy (Fig 3E).

The cilia immune-capture method was limited by low yield and further sample loss due to the instability of antibody-bound cilia, which hindered subsequent composition analysis by immunofluorescence or biochemistry. Therefore, we used a complementary method to increase yield, in which culture media were subjected to a series of filtration and centrifugation steps to concentrate ciliary material approximately 500-fold (Fig. 3F-G, S4B). As a positive control, serum-starved cells were artificially deciliated with high-calcium buffer (62) (see *Materials and Methods*). Immunostaining for axoneme markers (IFT88, α-tubulin, and acetylated tubulin) demonstrated increased abundance of ciliary structures, many with the dimensions (1.5-7 μm) expected of whole cilia (Fig. 3F). Immunoblotting against IFT88, α-tubulin, and acetylated tubulin further confirmed enrichment of ciliary proteins in the concentrated medium (Fig. 3G). These results indicate that whole cilia, including membrane and axoneme components, are shed during ciliary disassembly.

Together, these two independent ciliary isolation methods suggest that deciliation occurs during serum-induced ciliary disassembly, and is the phenomenon likely underlying *instant* ciliary disassembly dynamics.

### p60 katanin overexpression promotes *instant* ciliary loss

We next sought to gain insight into the regulation of deciliation. Although this process has been observed in several organisms (49,63–65), the underlying machinery and mechanism of deciliation have been elusive. A central requirement for complete deciliation is the disruption of axonemal microtubules at the ciliary base (44,51,66,67). The microtubule-severing enzyme katanin has been proposed as a candidate for this function in *Chlamydomonas* and *Tetrahymena*, and has also been shown to have a general regulatory relationship with primary cilia (46,51,66,68–71). Katanin is also required for microtubule rearrangement in mitotic spindle formation (69,70,72,73), but its role in primary cilia disassembly is poorly understood.

To examine the potential regulation of ciliary disassembly by katanin, we manipulated katanin levels by depletion or overexpression. We found that functional depletion of the catalytic subunit of katanin, p60 (KATNA1), by siRNA or expression of a catalytically inactive mutant (74–76) caused inviability in IMCD3 cells (data not shown), consistent with its function in mitosis. We then overexpressed p60 katanin by creating IMCD3 cells stably co-expressing SSTR3-GFP and turboRFP-p60. Consistent with previous reports (46,70,74,75,77,78), we observed diffuse localization of p60 in the cytoplasm in both tRFP- and tRFP-p60-expressing cells. In addition, in about 30% of ciliated cells, p60 localized near the base of the primary cilium (Fig. 4A). Cytoplasmic acetylated tubulin intensity was reduced in serum-starved and serum-stimulated tRFP-p60 cells compared to the control (Fig. S6A-B), consistent with increased severing and destabilization of microtubules caused by katanin overexpression (71,74,79). We conclude that overexpressed katanin is functional.

**Fig 4.**
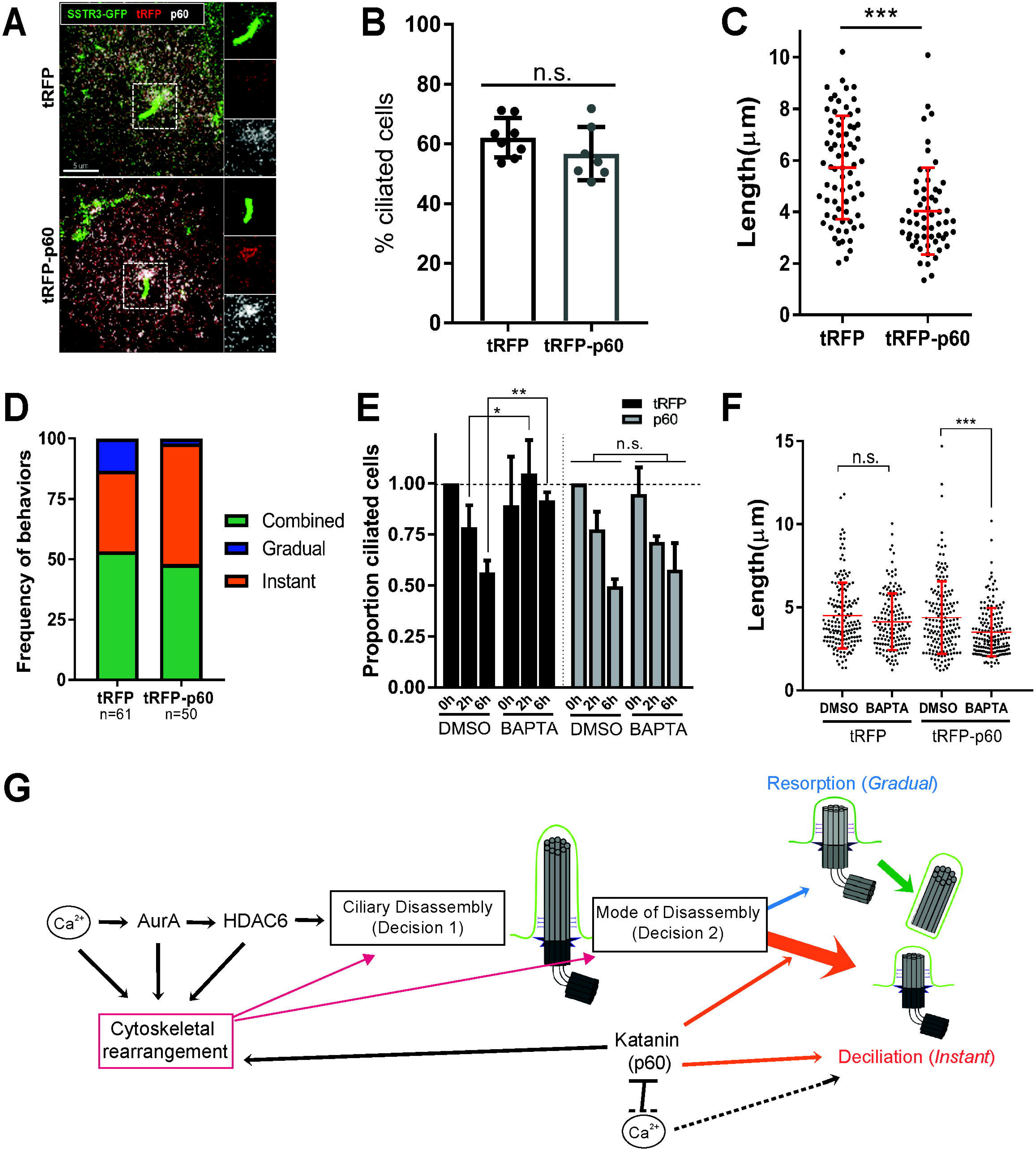
tRFP-p60 overexpression reduces ciliary length, promotes *instant* disassembly, and counteracts effects of [Ca^2+^] modulation on cilia. A-D) IMCD3-SSTR3::GFP cells stably expressing turboRFP (tRFP) or tRFP-p60 fusion. A) Serum starved tRFP and p60 cells were fixed, and stained with a polyclonal antibody against p60. (green – SSTR3::GFP; red – tRFP; white – total p60 stain). B) Percent ciliation calculated from fixed starved tRFP and tRFP-p60 cells. Data from at least 7 independent experiments. C) Cilia lengths measured from confocal stacks of live cells. Data pooled from at least 3 independent experiments, at least 50 cilia per experiment. Statistical significance from Mann-Whitney U test. D) Relative frequencies of *gradual*, *instant*, and *combined* dynamic behaviors in tRFP and tRFP-p60 cells. Data pooled from at least 4 independent experiments. E-F) Effects of BAPTA-AM on (E) ciliary abundance, disassembly and (F) length. Cilia counts from fixed populations, normalized to respective DMSO controls; length from live cells. tRFP and p60 cells were starved and pre-treated with DMSO, or 1 μM BAPTA-AM (30 min). E) BAPTA-AM pre-treated cells were subjected to serum stimulation and fixed at indicated time points. Data from 3 independent experiments, at least 100 cells counted per condition. Statistical significance was determined by unpaired t-test. G) Model for regulation of ciliary disassembly. Major regulators of ciliary disassembly function at two decision points – whether (Decision 1) and how (Decision 2) to disassemble the cilium, which is dominated by deciliation (orange), but also features resorption (blue), and combined resorption-deciliation (green). Katanin (p60) and Ca^2+^ can promote deciliation at Decision 2 independently and/or antagonistically. Regulators of disassembly may further contribute to either decision indirectly via modulation of cytoskeletal rearrangement (magenta). See text for full description.

Next, we determined whether tRFP-p60 katanin expression influenced overall assembly and disassembly of cilia. Total levels of ciliation in response to serum starvation (Fig. 4B) and stimulation (Fig. S5C) were unaffected, however, ciliary length was significantly reduced in tRFP-p60 cells (Fig. 4C). Furthermore, the extent of inhibition of ciliary disassembly via tubacin (19,24,34,80) and cytochalasin D (4,81,82) was unaffected in tRFP-p60 cells (Fig. S5C). Therefore, tRFP-p60 katanin expression influences ciliary structure, but is not sufficient to induce ciliary disassembly in serum-starved cells, and any effect of expression likely occurs downstream of HDAC6 activity.

We next asked whether tRFP-p60 expression affects ciliary disassembly dynamics in serum-stimulated cells. In cells expressing only tRFP, terminal cilium loss by *instant* dynamics comprised 86.7% of disassembly events (n=61, 33.3% *instant* and 53.3% *combined*), consistent with our analysis in Figure 2G. However, in tRFP-p60-expressing cells, *gradual* disassembly was virtually eliminated, while the frequency of *instant* disassembly increased, resulting in 98% of disassembly events featuring *instant* terminal cilium loss (n=50, 50% *instant* and 48% *combined*) (Fig. 4D). Thus, tRFP-p60 expression shifts the distribution of disassembly behaviors toward *instant* ciliary loss dynamics. These results suggest that increased katanin activity is capable of modulating ciliary disassembly behavior by promoting deciliation with *instant* dynamics, and that the distribution of heterogeneous disassembly behaviors is tunable by mechanistic regulators.

### Katanin p60 overexpression blocks the effects of Ca^2+^ modulation on ciliary disassembly

High intracellular calcium concentration ([Ca^2+^]_i_) can trigger deciliation, and many methods of experimentally inducing deciliation do so by raising [Ca^2+^]_i_ ((46,47,49–51,61,62,83–88), also see Figs. 3F-G & S3). p60-katanin function and Ca^2+^ are both necessary for experimentally-induced deciliation in isolated flagella (46). Additionally, calcium-calmodulin signaling functions upstream of AurA, a key regulator of ciliary disassembly upstream of HDAC6 (23,35,89), suggesting a broader role for cytoplasmic calcium in regulating ciliary disassembly via all of the behaviors we have identified. Thus, we hypothesized that calcium and katanin functions may cooperate in serum-induced ciliary disassembly. We asked whether, in the presence of high [Ca^2+^]_i_, the frequency of cilium loss would be exacerbated by expression of tRFP-p60. Cells were serum-starved to promote ciliation, and then treated with small molecule drugs to modulate [Ca^2+^]_i_ levels. In both tRFP- and tRFP-p60-expressing cells, raising [Ca^2+^]_i_ with either 1 μM ionomycin (90) or 190 μM dibucaine, reduced ciliary abundance and length to a similar degree (Fig. S5). Treating cells with 5 μM thapsigargin, which raises [Ca^2+^]_i_ by releasing ER stores (42,65, 67), also reduced ciliation but not cilia length (50) in control tRFP-expressing cells. Intriguingly, this effect was reversed in tRFP-p60-expressing cells, which exhibited a slight increase in ciliation and cilia length in response to thapsigargin (Fig. S5C-D). Overall, tRFP-p60-expressing cells did not exhibit elevated rates of ciliary loss induced by high [Ca^2+^]_i_.

Conversely, we asked what effects depletion of [Ca^2+^]_i_ would have on ciliary disassembly in tRFP- and tRFP-p60-expressing cells. BAPTA-AM, a cell-permeable Ca^2+^ chelator (92), did not affect ciliary length in starved cells, but inhibited ciliary disassembly in serum-stimulated tRFP cells, (Fig. 4E-F), consistently with published work (23). However, disassembly was not impaired in serum-stimulated tRFP-p60 cells (Fig. 4E), and ciliary length was reduced in starved cells (Fig. 4F). Therefore, tRFP-p60 katanin expression can overcome the requirement for [Ca^2+^]_i_ in ciliary disassembly. We summarize these results in a schematic in Fig. S6E. Overall, we found that overexpression of tRFP-p60 eliminated or reversed the effects of both increasing [Ca^2+^]_i_ with thapsigargin (Fig. S6C-D) and reducing [Ca^2+^]_i_ with BAPTA-AM (Fig. 4E-F). Taken together, these results indicate that [Ca^2+^]_i_ and katanin do not act cooperatively to promote ciliary loss. Rather, the negative regulation of cilia by [Ca^2+^]_i_ appears to be mitigated or reversed in the presence of excess p60 katanin.

## Discussion

We comprehensively characterized the disassembly dynamics and behaviors of IMCD3 primary cilia, a process critical to tissue homeostasis and development in vertebrates. Here we will first present a model for ciliary disassembly based on these data, and then discuss implications of that model.

Our results suggest that there are at least two mechanisms of ciliary disassembly in mammalian cells: resorption, in which the axoneme is depolymerized and ciliary contents are incorporated into the cell, and deciliation, in which the axoneme is excised near its base. We have incorporated these in a model (Fig. 4G) in which ciliary disassembly requires at least two major decision points – Decision 1: when to disassemble the cilium; and Decision 2: when to invoke instant deciliation during disassembly. Decision 1 is controlled by Aurora A, HDAC6, and other elements (24,34,54,80,93). Decision 2, in IMCD3 cells, results in deciliation immediately (*instant*), after a delay (*combined*), or not at all (*gradual*). We assume here that resorption of the cilium, as occurs in *combined* and *gradual* events, is the default mode of disassembly, and that rapid deciliation, when invoked, overrides the slower-acting resorption. Thus, cilium disassembly is a tunable decision in at least two ways – when to remove the cilium, and the mechanism by which that removal is accomplished, both of which are likely to differ in different cell types and different contexts.

We manipulated the system in several ways and found that the activity of overexpressed katanin biases disassembly events nearly exclusively toward deciliation, likely representing an intervention at Decision 2. Whether and how katanin might sever axoneme microtubules in deciliation, or whether it is indeed required for deciliation (94) remain critical outstanding questions. Katanin may also indirectly influence cilia, perhaps by severing centrosomal microtubules and interfering with ciliary protein trafficking (in line with a diffuse staining pattern at the cilium base, Fig. 4A), or by modulating the available pool of cytoplasmic tubulin (95,96). These hypotheses could help explain why tRFP-p60 expression negatively affects cilia length but not formation.

Calcium is necessary for ciliary disassembly (23) and sufficient to drive deciliation (63,64,85,97), likely acting at both Decisions 1 and 2. Although the role of calcium is likely multifaceted, it seems that calcium and increased p60 katanin activity may function independently, and also negatively interact at the Decision 2 nexus (Fig. 4E-F, S6C-E). Despite the requirement for Ca^2+^ for disassembly (Decision 1) and deciliation (Decision 2) in normal conditions, tRFP-p60 cells were able to undergo ciliary disassembly after [Ca^2+^]_i_ chelation, likely via deciliation. This suggests the existence of a Ca^2+^-independent deciliation pathway, as has been suggested previously (64). Alternatively, overexpressed p60 may promote disassembly via resorption or other means in the absence of [Ca^2+^]_i_. This relationship may be consistent with the finding that calcium binding inactivates p60 severing activity *in vitro* (76). We speculate that the activities of katanin and [Ca^2+^]_i_ in modulating ciliary behavior may depend on their relative levels in the cell (Fig. S6E).

We emphasize that disassembly of the cilium, a major organelle of the cell, likely involves a complex interplay between general ciliary trafficking and regulation, cytoskeletal dynamics, and intracellular signaling ((4,9), Fig. 4G), and that our manipulations of calcium and katanin are best viewed as initial probing into the molecular nature of these decisions. Further work will be required to understand the molecular mechanisms and must take into account that many key regulators of ciliary disassembly have additional roles in cytoskeletal regulation (22,32,54) and other cellular functions, including calcium (98,99), AurA (100–102), HDAC6 (4,103–106), and katanin (69,73,107).

Ciliary disassembly behaviors varied both in the dynamics (10^−3^ to 10^1^ μm/min) and physical process (resorption vs. deciliation). The co-existence of resorption and deciliation in the same cilium (*combined* disassembly) is intriguing and may suggest independent or differential regulation of the distal and proximal portions of the cilium (36). It might be, for example, that structural features of the axoneme, such as doublet-singlet microtubule interface or post-translational modifications, contribute to differential regulation of ciliary regions (108– 110). We were unable to observe the axoneme directly in our live cell imaging experiments, however, in fixed serum-stimulated cells, we identified several non-canonical structures that hint at the fate of the axoneme during ciliary disassembly. These included discontinuous axoneme staining that might reflect partial breaks away from the base, short axoneme stubs that could represent a remnant of a severed or resorbed cilium (111–113), and even axonemes without corresponding ciliary membrane staining, possibly representing a portion of axoneme retracted into the cell, as has been previously reported (39). While these interpretations are speculative due to the markers used and the nature of static representations of this dynamic process, the relatively low abundance of these structures in starved and tubacin-treated conditions indicates that they may represent ciliary disassembly intermediates.

We observed deciliation as a means for cell cycle-linked ciliary disassembly, whereas previous descriptions of such behavior in mammalian cells have only been under conditions of experimentally induced stress (50,52). The morphology and composition of isolated cilia confirmed two major points supporting the interpretation that the observed *instant* disassembly events represent deciliation: 1) the entire ciliary membrane can be shed from cells as an intact structure; and 2) shed ciliary membranes contain tubulin, suggesting that the axoneme is severed and shed along with the ciliary membrane. Interestingly, release of a small membrane segment from the ciliary tip, referred to as apical abscission, decapitation, or release of ciliary ectosomes, has been described in several contexts (9,80,114–117), but it is unclear whether axoneme components are present in these structures. Ciliary decapitation was previously reported as an initiating step in the cilia disassembly process (80), as well as in assembling cilia (9). Although we observed similar tip-shedding behaviors in IMCD3 cells (M. Mirvis, unpublished results), their relationship to ciliary disassembly was not clear and they did not necessarily precede disassembly events. It would be particularly interesting to test whether there is a mechanistic link between these events by investigating whether PI(4,5)P_2_- and F-actin-mediated membrane constriction that drives decapitation (80) participates in deciliation from the ciliary base as well.Finally, the implications of our findings for understanding cilia disassembly should be considered in the context of the general features of cultured kidney-derived epithelial cells, in comparison to and contrast with cilia in other experimental approaches, cell types, tissues, and organisms. Although most cilia share a similar core ciliary structure and associated machinery, differences in structure and function (i.e. primary vs. motile/specialized) and the relative size of cilia to the cell body, are likely to be relevant to the mechanisms of ciliary assembly and disassembly.

## Acknowledgements

We thank Jonathan Indig for critical assistance with developing and writing the Matlab algorithm, Fan Ye and the Max Nachury lab for the gift of IMCD3-SSTR3::GFP cells, Martijn Gloerich for assistance with lentivirus, Daniel Cohen and Caitlin Collins for technical assistance with cilia immune-capture, Lydia Joubert and the Beckman Cell Sciences Imaging Facility for assistance with SEM sample preparation and imaging. We thank Jessica Feldman, Lucy O’Brien, and members of the Nelson and Stearns laboratories for invaluable discussion of the methods and results; Ljiljana Milenkovic, Jennifer Wang, and Jackson Liang for thoughtful feedback on the manuscript. Research reported in this publication was supported by the National Institute of General Medical Sciences of the National Institutes of Health under award number T32GM007276 (M.M.), R35 GM118064 to W.J.N. and R01GM121424 to T.S. The content is solely the responsibility of the authors and does not necessarily represent the official view of the National Institutes of Health.

## Materials & Methods

### Cell Culture

IMCD3 cells were grown in DMEM-F12 medium with 10% fetal bovine serum and 1% penicillin-streptomycin-kanamycin antibiotic cocktail. Cells were passaged every 2-3 days at a dilution of 1:10-1:20. Cells were tested for mycoplasma with Sigma LookOut Mycoplasma PCR Detection Kit (Cat#MP0035) as directed by the manufacturer, and incidences of mycoplasma contamination was treated with Mycoplasma Removal Agent (MP Biomedicals,#093050044). Following decontamination, experiments that were potentially affected by mycoplasma contamination were repeated at least three times to determine any difference in results, and no significant differences were observed.

### Serum Starvation and Stimulation

Cells were seeded in 24- or 6-well dishes with glass coverslips for imaging following fixation, or 35 mm glass-bottomed MatTek dishes (#P35G-0-10-C) for live imaging. 24-well dishes were seeded at a density of 1.5×10^4^ cells and 6-well and 35 mm MatTek dishes at 1-1.5×10^5^, to achieve 50-70% confluence next day. For serum-starvation, cells were washed once with 0.2% DMEM-F12 + PSK, then grown in 0.2% DMEM-F12 + PSK for 24 hr. Serum stimulation was by either re-addition of FBS directly to dishes to 10% final concentration, or replacement with 10% FBS DMEM-F12.

### Antibodies

The following antibodies and dilutions were used. Acetylated tubulin mouse monoclonal 6-11B-1 (1:1000 for IF & WB) (Sigma-Aldrich Cat# T7451); Pericentrin rabbit polyclonal Poly19237 (1:500 for IF) (Covance Cat #PRB-432C, now BioLegend); Arl13b rabbit polyclonal (1:250-1:500 for IF) (Proteintech Cat# 17711-1-AP); N19-SSTR3 antibody (rabbit polyclonal) (Santa Cruz Cat #sc-11610, discontinued); IFT88 rabbit (1:500 for IF & WB) (GeneTex, Cat#79169); α-tubulin YL1/2 (1:1000 for IF & WB), (ThermoFisher #MA1-080017); alpha-tubulin DM1a (1:1000 for IF & WB) (ThermoFisher #62204); rabbit monoclonal anti-p60 EPR5071, (1:250 IF), (Abcam Cat# ab111881); rabbit polyclonal anti-KATNA1, (1:100-250 IF) (Proteintech Cat#17560-1-AP). Anti-rabbit GFP (1:250) (Life Technologies, #A11122); Anti-mouse GFP (1:1000) (Roche, #11063100). Secondary antibodies used were: Anti-mouse Rhodamine (1:1000) (Jackson ImmunoResearch, #715-295-150), Anti-rabbit FITC (1:1000) (Jackson ImmunoResearch, #111-095-003), Anti-rabbit Alexa647 (1:200) (Life Technologies, #A21245), Anti-mouse Alexa647 (1:200) (Life Technologies, #A21236), Hoescht (1:1000-2000) (Molecular Probes, #H-3570).

### Chemicals

Tubacin (Sigma-Aldrich, #SML0065) at 2 μM in DMSO; dibucaine hydrochloride, (Sigma-Aldrich #285552) at 190 μM in DMSO. The following were used at 1μM in DMSO: Cytochalasin D (Sigma-Aldrich #PHZ1063), Thapsigargin, (Sigma-Aldrich #T9033). BAPTA-AM (Sigma-Aldrich #A1076). Ionomycin, 10 mM stock was a gift from the Rich Lewis laboratory, Stanford Univ.

### Generation of stable cell lines

IMCD3-SSTR3::GFP-mCherry::PACT: mCherry::PACT was cloned from a pLV plasmid (pTS3488, created by multi-site Gateway cloning by Christian Hoerner) onto pLV-Puro-EF1a construct using Gibson cloning. Lentivirus with the cloned construct was generated in HEK293T and used to infect IMCD3-SSTR3::GFP (gift from Nachury laboratory, (55)) under selection with 800 mg/mL puromycin for 4-5 days. Infected cells were FACS-sorted into polyclonal populations by mCherry fluorescence intensity, and a pool of low-expressing cells was selected to prevent over-expression phenotypes of a centrosomal protein.

#### Katanin expression constructs

Mammalian expression constructs for turboRFP and turboRFP::p60 (p60 domain of mouse katanin) were designed and ordered from VectorBuilder. All constructs were amplified by transformation in DH5α and maxi-prep (Qiagen #12165). IMCD3-SSTR3::GFP cells were transfected with each construct with ThermoFisher Lipofectamine 3000 according to manufacturer's protocol (#L3000015). The next day, cells were subjected to G418 selection (800ng/μL for 5-6 days). Cells were sorted by FACS into low-, medium-, and high-expressing pools, and maintained in DMEM-F12 10% FBS + PSK and 250 ng/μL G418 to maintain transgene expression.

### Transient transfection

mCherry-α-tubulin mammalian expression construct (gift from Angela Barth) was transfected into IMCD3-SSTR3::GFP cells. Transfections were performed using Lipofectamine 3000 transfection reagent according to manufacturer’s protocol.

### Immunofluorescence microscopy

Generally, fixation for immunofluorescence microscopy was done with 100% methanol for 5 minutes at -20°C, followed by washes with 0.1% Triton X-100 in PBS at room temperature for 2 minutes, and washed 3 times in PBS. Samples were blocked for 1 hr at RT°C or overnight at 4°C in 2% BSA, 1% goat serum, 75mM NaN_3_. Antibodies were diluted to the indicated concentrations in blocking buffer. Primary antibody incubations were performed for 1 hr at RT°C or overnight at 4°C. Secondary antibody incubations were performed for 1-2 hr at RT°C. Following each antibody incubation, samples were washed 3 times in PBS + 0.05% Tween-20 for 5 mins each at RT°C.

Images were acquired with a Zeiss Axiovert 200 inverted epifluorescence microscope and a 63× objective, or a Leica SP8 scanning laser confocal microscope with LASX Software, using mercury or argon lamps with white light laser excitation, and a 63× 1.4 NA oil objective. Exposure times were constant during each experiment. For imaging of serum-starved and serum-stimulated cells, fields of view were selected based on DAPI staining by two critieria: 1) to select for moderate cell density, in order to avoid effects of high density on cell cycle and ciliation; and 2) to eliminate bias in % cilia quantifications from scanning by ciliary markers.

### Live-cell confocal microscopy

Cells were cultured on glass-bottomed Mattek dishes and imaged in DMEM-F12 media with 15mM HEPES without phenol red. Movies were acquired 4-12 hr after serum stimulation with a Leica SP8 scanning laser confocal microscope using 0.5 μm z-slices, 30-90 sec intervals, autofocus, in a 37°C incubator, and red and green channels were acquired simultaneously. The video file was saved as .lif from LASX software and opened in Imaris x64 8.0.2 as a 3D render for analysis of cilia disassembly dynamics and basal body positioning.

### Data analysis

Cilia counts and length measurements were performed either manually in Fiji or Imaris x64 8.0.2 and 9.2.1, or through semi-automated detection in Imaris. Manual analysis involved detecting ciliary membrane, marked by an enrichment of SSTR3::GFP above background threshold, that were adjacent to a centriole (mCherry::PACT in dual-fluorescent cells or pericentrin immunofluorescence in single-(SSTR3::GFP-expressing) or non-fluorescent cells), to distinguish from accumulations of SSTR3+ membrane elsewhere in the cell. Manual length measurements in Fiji were made with the line function, and in Imaris with the Measurement tool. Generally, single z-plane images were analyzed in Fiji or Imaris, while confocal z-stacks were analyzed in Imaris which allowed more accurate length measurement due to the 3D render (Surpass) capability. When possible, length measurements in confocal images were semi-automated in Imaris using the Surfaces function to create an artificial object encompassing the ciliary membrane, and exporting Bounding Box data as a proxy for length (the longest dimension of the object).

For live cell serum stimulation experiments, movies were visually scanned in Imaris for examples of disassembling cilia. Images of each disassembling cilium were cropped by time (from t0 to several mins after complete loss), and position (restricted to area of occupancy during the that time window), and then saved in a separate file. To generate ciliary length curves, the ciliary membrane was isolated as an artificial object using the Surface function. When possible, the object was automatically tracked over consecutive time points with length data generated at each time point. In cases where automatic tracking was not possible due to low signal-to-noise of ciliary membrane fluorescence, measurements were taken manually at 15-30 minute intervals until the initiation of ciliary disassembly, and at each time point during the disassembly event.

#### Matlab

Raw length measurement data from disassembling cilia movies were compiled in to an Excel spreadsheet. A Matlab algorithm imported the data, performed smoothing and calculations of cilium start point, and generated an output file containing disassembly rates, start and end time, start and end length, and proportion length lost per disassembly stage. Algorithm strategy is described in the text and Supplement.

### Cilia Isolation

Cell culture: clones of IMCD3 cells, either an unsorted stably expressing GFP-SSTR3 or FACS sorted for medium expression of GFP-SSTR3, were grown on 15cm dishes at 3×10^6^ cells/dish in DMEM/F12 with 10% FBS and antibiotics for 24 hours. The cells were washed 3× with HDF wash buffer, and media was replaced with DMEM/F12 and 0.2% FBS and antibiotics (serum-starved) for 24 hours. Then all dishes were washed 3× with HDF buffer and half received phenol-red free DMEM/F12 with 0.2% FBS (serum-starved), and the other half received phenol red free DMEM/F12 with 10% FBS (serum-stimulated) for 24 hours. Total serum starved time was 48 hours and total serum stimulated time was 24 hours.

#### Immune-capture Method

Preparation of antibody-immobilized imaging dishes: Glass in 35 mm glass-bottomed imaging dishes (MatTek) was functionalized by plasma cleaning at 250 moor, Low setting, 45-60 seconds. Dishes were silanized with 500 μL of 2.5% triethoxysilyl-undecanal (TESU) in 100% ethanol, covered with Parafilm and incubated at RT°C for 1hr. Dishes were washed 3× with 100% ethanol, then baked at 85°C for 3 hrs. Next, silanized dishes were treated with the following series of reagents for 1hr at RT°C unless stated otherwise, with 3 PBS washes in between steps: 1) 50mM NHS-LC-LC-biotin in water, 2) 5mg/ml neutravidin for 1hr at RT°C, 3) 300 μg/ml biotin-Protein A 4) block with 15mM D-biotin in DMSO for 30 min, 5) anti-rabbit SSTR3 N19 (extracellular N-terminus) antibody (100-200 μg/mL) at 37°C, followed by one PBS wash. These protocols adapted from Dr. Nicholas Borghi (118).

#### Sample preparation

Culture medium was collected and subjected to centrifugation for 10 min at 1000xg at 4°C to remove large cell debris. Samples were then kept on ice until plating on treated dishes or stored at 4°C for a maximum of 1 day. 4mL serum-stimulated or -starved medium was incubated on a treated MatTek dish overnight at 4°C, followed by 3 gentle PBS washes. Samples were then imaged directly, without fixation with a Leica SP8 confocal microscope.

#### SEM

Antibody-immobilized MatTek dishes incubated with serum stimulation medium were fixed for SEM in 4% PFA, 2% glutaraldehyde, and 0.1M Na cacodylate. Glass bottoms were removed, processed for imaging, and imaged with a Hitachi S-3400N VP SEM scope in the Beckman Imaging Facility, Stanford Univ.

#### Filter-Spin Concentration Method

Harvest of Cilia: Deciliation of starved IMCD3 cells (positive control): Serum-stimulated or -starved culture medium, or fresh culture medium (with 10% FBS, an additional control) was removed from 6 150cm dishes, combined and centrifuged at 200xg at 4°C for 5 mins in the A-4-81 rotor, Eppendorf 5810R centrifuge. Cells were washed 2x with warm PBS containing 0.4% EDTA. 10 mL was added to a MatTek dish and incubated for 10 min at 37 °C, followed by gentle up and down pipetting to remove cells from dish. An aliquot of cell suspension was removed for cell count. Cells were centrifuged at 13000 x g for 5 mins at RT°C. The cell pellet was resuspended in 5 mL ice cold deciliation buffer (62) (112 mM NaCl, 3.4 mM KCl,10 mM CaCl_2_, 2.4 mM NaHcO_3_, 2 mM HEPES, pH7.0 and a protease inhibitor tablet [Roche]). The cell suspension was incubated at 4 °C for 15 mins with rigorous end-over-end rotation, and then centrifuged at 1000 x g for 5 mins at 4 °C in an Eppendorf centrifuge. The resulting supernatant was used for biochemistry and immunostaining.

#### Biochemistry

Half of the supernatant material from deciliated, serum starved or -stimulated cells was centrifuged at 21,000xg for 15 min in JA25.5 rotor in Beckman Coulter Avanti J-25I centrifuge at 4°C. The supernatant was carefully removed, and pellets were resuspended in 160 μl of sample buffer (1% SDS, 10mM Tris-HCl, pH 7.5, 2mM EDTA). Samples were boiled at 95°C for 8 mins, and equal volumes were separated by 10% PAGE and transferred to PVDF. Blots were blocked (2% BSA, 1% normal donkey and goat serum in TBS, pH 7.4) for 1hr at RT°C or overnight at 4°C. Membranes were blotted with YL1/2 (1:1000), mouse acetylated-tubulin antibody (1:1000), and IFT88 rabbit antibody (1:500) in blocking buffer for 1 hr at RT°C. Blots were washed 5x with TBST. Secondary anti-rabbit, anti-mouse, or anti-rat antibodies labeled with either Alexa Fluor 680 (InVitrogen, #A21058) or IRDye800CW (Li-Cor Biosciences, #926-32213), at 1:30,000 dilution were incubated with blots for 30 min at RT. Blots were washed 5x with TBST and scanned on Licor Odyssey scanner (Li-Cor BioSciences).

#### Immunofluorescence

Half of the supernatant material from deciliated, serum-starved, or -starved and -stimulated cells was concentrated using a 250ml 0.2μm PES filter unit with house vacuum to reduce the volume to 2 mL, and finally a Millipore Ultrafree-MC filters (PVDF 0.2μm size #UFC30GV100) to reduce the volume to ~0.5 mL. 5 μl of concentrated supernatant was pipetted onto an acid-treated glass slide. A 22 mm acid-treated circular glass coverslip was placed on the sample, and the slide was immediately plunged into liquid nitrogen for ~5 sec. After removing the slide, the coverslip was removed and fixed in -20 °C 100% methanol for 5 mins. Immunofluorescence staining was performed as described above.

### Statistics

All analyses were performed in GraphPad Prism. Statistical tests used for each analysis are indicated in the Figure legends. No explicit power analysis was used to determine sample size. All experiments were performed with at least three biological replicates, i.e. samples from independent cell culture passages. When used, technical replicates (i.e. repeats from the same cell culture passage) were averaged for each biological replicate. In brief, comparisons of mean values such as mean percent cilia across replicate experiments were compared using an unpaired t-test. Analyses of individual measurements such as cilia length were subjected to normality tests (Kolmogorov-Smirnoff, D’Agostino & Pearson, and Shapiro-Wilk). If data passed all normality tests, unpaired t-test was used, if not the Mann-Whitney U test was used. If data passed normality by some tests but not others, both types of analyses were performed. Results were similar between parametric and nonparamentric tests unless stated otherwise.

## Supporting Material

**Figure S1. Characterization of immunostained serum stimulated cells.** A) Serum starved and stimulated cells were immunostained for pericentrin to mark the basal body (PCNT, white) and acetylated tubulin to mark the axoneme (acTub, red). Scale bar, 10 μm. B) Serum stimulation is accompanied by an increase in mitotic cells, inhibited by tubacin treatment. Quantifications are based on means of 3 independent experiments with 150-200 cells analyzed per condition per replicate. Error bars – S.E.M.

**Figure S2. Schematized workflow for automated analysis of ciliary disassembly dynamics**. A)10 control non-disassembling cilia from starved cells were imaged at 90-sec intervals over a 12 hr period and analyzed. Cumulative metrics are shown here (μm). Slight changes in length and stochastic length fluctuations are inherently present, and used as a baseline for analysis of disassembling cilia. Marked in yellow – maximum standard deviation of length is used as a proxy for length measurement error, and is thus used as a threshold length for instant disassembly; average slope of best fit line represents background decline in length over 12 hr imaging period. B) Matlab workflow. Scale bar, 5 μm. Raw data are run through a smoothing function. Derivatives are calculated, then normalized to background reduction (A) to identify start point of disassembly event. Lastly, disassembly behaviors (gradual, instant, and combined) are assigned as described in the text.

**Figure S3. Dynamics of dibucaine-induced ciliary shedding are consistent with serum-induced instant disassembly**. Starved cells were treated with 190 μm dibucaine and imaged by confocal microscopy at 30 sec intervals. A) Still images from a representative ciliary shedding show complete ciliary loss in under 30 sec. B) Length measurements from A) show instant disassembly dynamics.

**Figure S4. Schematics of cilia capture methods.** A) Immune-capture of shed cilia. Medium from serum stimulated cells is incubated on an imaging dish bearing immobilized antibody against the SSTR3 membrane marker. B) Filter-spin concentration of shed cilia. Medium from serum-stimulated cells is concentrated either by centrifugation pelleting for subsequent Western blot or by vacuum & centrifugation filtration for immunofluorescence. Medium from cells subjected to artificial deciliation was included as a positive control.

**Figure S5. tRFP-p60 overexpression reduces cytoplasmic acetylated tubulin intensity, but does not impair overall serum-induced ciliary disassembly**. A) Starved and 2 hr-stimulated tRFP and tRFP-p60 cells were fixed and stained for acetylated tubulin (red, center column). Scale bar, 20 μm. Insets show acet. tubulin channel alone. Inset scale bar, 10 μm. B) Quantification of A). Mean cellular acTub intensity was normalized to cellular tRFP intensity. Data pooled from 3 independent experiments, 80-100 cells per condition. Statistical significance from Mann-Whitney U test. C) D) Serum starved cells pre-treated with DMSO, 2 μM Tubacin, or 1μM CytoD were serum stimulated (6 hr). Proportion of ciliated cells was calculated by normalizing to Starved + DMSO for each cell line (dotted line). N = 3 experiments.

**Figure S6. tRFP and tRFP-p60 cells exhibit similar responses to elevated [Ca^2+^]_i_ by ionomycin and dibucaine, but not to thapsigargin**. tRFP and tRFP-p60 cells were starved and pre-treated with DMSO, A-B) 190 μM dibucaine (30m), 1 μM ionomycin (30m), and C-D) 5 μM thapsigargin (Thps) (1 hr). Cilia counts are from fixed populations and normalized to respective DMSO controls. Length measurements are from confocal z-stacks of live cells. Data from 3 independent experiments, at least 100 cells counted per condition. Statistical significance was determined by unpaired t-test. E). Model summarizing results from Fig. 4E-F & S6C-D. Orange bar – tRFP-p60 cells have no defect in overall ciliary disassembly, but undergo *instant* deciliation more frequently. We make several assumptions to approximate the relative differences in p60 and [Ca^2+^]_i_ levels between our experimental manipulations: 1. tRFP-p60 cells have “*excess*” p60 compared to “*normal*” levels in tRFP control cells. 2. Thapsigargin may produce a [Ca^2+^]_i_ that is physiological, but higher than the ground state (89) (labeled “*excess*” on the figure), whereas dibucaine and ionomycin induce “*high excess*” levels of [Ca^2+^]_i_.

## Supplementary Videos

**Video S1.** *Primary cilium of a serum starved cell*. Imaged at 90 second intervals over 12 hrs, 45fps. The cilium undergoes rapid length fluctuations and a slight overall reduction in length at 0.003 μm/min. See Fig. 2A.1.

**Video S2**. *Primary cilium disassembling by gradual dynamics*. Imaged at 90 second intervals, 45fps. Cilium disassembles with an overall rate of 0.016μm/min. See Fig. 2A.2.

**Video S3**. *Primary cilium disassembling by instant dynamics*. Imaged at 90 second intervals, 15fps. Cilium is lost in under 90 seconds, approximate minimum rate of 4.72μm/min. See Fig. 2A.3.

**Video S4**. *Primary cilium disassembling by combined dynamics*. Imaged at 46 second intervals, 25fps. Cilium undergoes gradual shortening (0.029μm/min), followed by instant loss (>5.69μm/min). See Fig. 2A.4.

**Video S5**. *Primary cilium shedding*. Imaged at 54 second intervals. Ciliary membrane is released from cell surface (27:12-29:00). See Fig. 3A.

